# Molecular changes in *Mesembryanthemum crystallinum* guard cells underlying the C_3_ to CAM transition

**DOI:** 10.1101/607333

**Authors:** Wenwen Kong, Mi-Jeong Yoo, Jerald D. Noble, Theresa M Kelley, Jing Li, Matias Kirst, Sarah M. Assmann, Sixue Chen

## Abstract

Crassulacean acid metabolism (CAM) is a specialized type of photosynthesis: stomata close during the day, enhancing water conservation, and open at night, allowing CO_2_ uptake. *Mesembryanthemum crystallinum* (common ice plant) is a facultative CAM species that can shift from C_3_ photosynthesis to CAM under salt or drought stresses. However, the molecular mechanisms underlying the stress induced transition from C_3_ to CAM remain unknown. Here we determined the transition time from C_3_ to CAM in *M. crystallinum* under salt stress. In parallel, single-cell-type transcriptomic profiling by 3’-mRNA sequencing was conducted in guard cells to determine the molecular changes in this key cell type during the transition. In total, 495 transcripts showed differential expression between control and salt-treated samples during the transition, including 285 known guard cell genes, seven CAM-related genes, 18 transcription factors, and 185 other genes previously not found to be expressed in guard cells. *PEPC1* and *PPCK1*, which encode key enzymes of CAM photosynthesis, were up-regulated in guard cells after seven days of salt treatment, indicating that guard cells themselves can transition from C_3_ to CAM. This study provides important information towards introducing CAM stomatal behavior into C_3_ crops to enhance water use efficiency.

**Summary statement:** We determined the timing of salt induced transition of common ice plant from C_3_ to CAM and identified transcriptomic changes during the transition. The data support the notion that guard cells themselves can transition from C_3_ to CAM.

## 1. INTRODUCTION

Global climate change is causing an increase in frequency of extreme drought and heat events and reducing fresh water and arable land for agriculture (Kaushal *et al*. 2017; Ziska *et al*. 2016). The impacts of extreme weather and drought are exacerbated by the demands of a growing human population, predicted to reach nine billion by 2050 (Borland *et al*. 2014). Therefore, improving the water-use efficiency (WUE) of agricultural crops is crucial to sustain productivity under increasing abiotic stresses and to expand cultivation to marginal lands, thereby enhancing food security.

In crassulacean acid metabolism (CAM), a specialized type of photosynthesis, stomata close during the day and open at night. CO_2_ is taken up at night and phosphoenolpyruvate (PEP) is converted to oxaloacetate by phosphoenolpyruvate carboxylase (PEPC). Oxaloacetate can be subsequently transformed into malate by malate dehydrogenase and transported into the vacuole. During the day, the organic acids are exported from the vacuoles and decarboxylated to produce PEP or pyruvate and release CO_2_ for light-driven carboxylation via ribulose-1,5-bisphosphate carboxylase/oxygenase (Rubisco) in the Calvin cycle (Owen & Griffiths 2013). The PEP is recycled as the substrate to assimilate CO_2_ during the night or used for synthesis of carbohydrates (Borland *et al*. 2014). By shifting atmospheric CO_2_ uptake to nighttime, when evapotranspiration rates are drastically reduced compared to the day, CAM plants achieve 3- to 6-times higher WUE than C_4_ and C_3_ plants (Nobel 1996). Therefore, introducing CAM into C_3_ crops could greatly improve WUE and drought tolerance of C_3_ plants.

*Mesembryanthemum crystallinum* (common ice plant) is a facultative CAM plant – it can shift between C_3_ and CAM. When *M. crystallinum* is grown under non-stress conditions, it can complete its life cycle solely with C_3_ photosynthesis (Adams *et al*. 1998). However, under stress conditions, such as water-deficit, salinity or high light, *M. crystallinum* can perform all the physiological features of CAM (Winter & Holtum 2005; Winter & Ziegler 1992). The shift from C_3_ to CAM in *M. crystallinum* may be mediated by a calcium-dependent signaling pathway. Pretreating leaves with a calcium chelator (ethyleneglycol-bis(aminoethyl ether)-N,N-tetraacetic acid) inhibits stress-induced transcription of *PEPC1*, NAD-glyceraldehyde-3-phosphate dehydrogenase gene 1 (*GapC1*) and cytosolic NAD-malate dehydrogenase gene 1 (*Mdh1*), which are all important for CAM (Taybi & Cushman 1999). In addition, increased activity of the antioxidative stress system (e.g., superoxide dismutase) caused by salinity and high irradiance (Hurst *et al*. 2004), or H_2_O_2_ applied to *M. crystallinum* roots (Surowka *et al*. 2016), can also induce the C_3_ to CAM transition. The inducibility of CAM and the biochemistry of C_3_ and CAM in the same cells (unlike C_4_ photosynthesis) make *M. crystallinum* an excellent system to study the mechanisms underlying the C_3_ to CAM transition.

Previous studies have revealed omics level changes in *M. crystallinum* in response to salt stress treatment. The first microarray study by Cushman *et al*. (2008) used five-week-old plants (as C_3_), and plants plus 14 days of salt stress (as CAM). A total of 1457 genes showed more than two-fold changes in mRNA steady-state levels between control (C_3_) and salt treatment (CAM). Many of the differentially regulated genes are involved in CAM-related C4 acid carboxylation/ decarboxylation, glycolysis/gluconeogenesis, starch metabolism, protein degradation, transcriptional activation, signaling, stress response, and transport (Cushman *et al*. 2008). Using next-generation sequencing, Tsukagoshi and co-workers (2015) identified 53,516 cDNA contigs from *M. crystallinum* roots and provided a transcriptome database (Tsukagoshi *et al*. 2015). They found that ABA responsive genes, a sodium transporter (*HKT1*), and peroxidase genes exhibited opposite responses to 140 mM NaCl treatment in *Arabidopsis* (C3) and ice plant (CAM) (Tsukagoshi *et al*. 2015). Oh and co-workers (2015) constructed a reference transcriptome containing 37,341 transcripts from control and salt-treated epidermal bladder cells (EBCs) of *M. crystallinum*, and 7% of the transcripts related to ion transport and signaling were salt stress responsive. At the small RNA level, Chiang *et al*. (2016) used roots of 3-day-old *M. crystallinum* seedlings and found 135 conserved miRNAs belonging to 21 families. The expression of mcr-miR159b and 166b, predicted to target transcription factors such as MYB domain protein 33, homeobox-leucine zipper family protein (HD-ZIP), and TCP4, were induced by salt treatment and mcr-miR319 expression was repressed. The proteome, ionome, and metabolome of *M. crystallinum* epidermal bladder cells have also been investigated (Barkla & Estrella 2015; Barkla *et al*. 2016).

The above studies identified numerous candidate genes, miRNAs, proteins and metabolites, which may inform efforts to improve plant salt tolerance. However, they have focused on either the steady-state levels of the C_3_ and/or CAM photosynthetic modes or the stress tolerance of *M. crystallinum*, not the transition from C_3_ to CAM. Therefore, the molecular mechanisms underlying this stress-induced transition remain unknown. Here we determined the critical transition time from C_3_ to CAM in *M. crystallinum* by measuring several key attributes, including titratable acidity, stomatal aperture, gas exchange, CAM-related enzyme activity, and CAM-related gene expression. Since reversed stomatal movement behavior is essential for CAM development, we tested the hypothesis that guard cells themselves undergo transition from C3 to CAM using single-cell-type transcriptomics.

## 2. MATERIALS AND METHODS

### 2.1. Plant material and growth conditions

*Mesembryanthemum crystallinum* seeds were germinated on vermiculite moistened with 0.5× Hoagland’s solution (Hoagland & Arnon 1950). One-week old seedlings were transferred to 32 ounce containers. The plants were grown in a growth chamber under 200 *μ*mol m^−2^ s^−1^ white light with a 12h (26°C) day /12h (18°C) night cycle, and watered daily with 0.5× Hoagland’s solution with micronutrients (50 ml/per plant). Four-week-old plants were treated by irrigating with 0.5× Hoagland’s solution containing 500 mM NaCl, in which the salt was provided as a source of stress to induce CAM (Cushman *et al*. 2008). Control plants were continuously watered with 0.5× Hoagland’s solution. The third pairs of leaves from the control and salt-treated plants were collected on day 5, day 6 and day 7 following the initiation of the salt treatment. Unless stated otherwise, three biological replicates from three different sets of plants were used.

### 2.2. Leaf nocturnal acidification

Leaf titratable acidity was measured as previously described (Cushman *et al*. 2008) with minor modifications. Leaves were harvested from control and salt-treated plants at the end of the night period (8 am). Fresh weight of the collected leaves was measured, followed by snap freezing in liquid N_2_ and storage at −80°C until analysis. Frozen leaves were ground to a fine powder using a mortar and pestle, followed by further homogenization in 80% methanol at a ratio of 7.5 ml/gram of fresh weight at room temperature. The homogenate was allowed to warm to room temperature. Aliquots of the methanol extracts were titrated against 5 mM NaOH to a neutral endpoint (pH = 7.0), using a pH meter. Leaf titratable acidities were expressed as μmol H^+^ g^−1^ fresh weight.

### 2.3. Net CO_2_ Exchange

Gas exchange measurements were performed at day 5, day 6 and day 7 of salt treatment. Net CO_2_ uptake was measured using a LI-6800 system (LI-COR Inc., Lincoln, NE, USA), and parameters were calculated using the manufacturer software. Conditions for measuring net CO_2_ uptake, stomatal conductance, and transpiration rates were: photon flux density 200 µmol m^−2^ s^−1^, chamber temperature 18°C/night and 26°C/day, flow rate 500 µmol s^−1^, relative humidity 50%, and 400 ppm CO_2_ reference. The chamber covered approximately 7 cm^2^ area of the ice plant leaf to measure net gas exchange. For each time point at least five plants were used. Rates of net CO_2_ uptake, stomatal conductance, and transpiration were retrieved from the LI-6800 data.

### 2.4. Stomatal aperture assay

The epidermis of control and salt-treated plant leaves were removed using 10 cm wide tape at 4 am and 4 pm, during the CAM transition process. After peeling, minor contamination from adhering mesophyll cells was removed by scratching the epidermis using a scalpel blade, and clean peels were directly used for imaging stomatal apertures. For stomatal aperture measurement, 60 – 80 stomata were randomly selected, and the sample identity was blinded during the measurement. Stomatal length and width were measured as described (Savvides *et al*. 2012), and stomatal aperture was estimated by the length/width ratio.

### 2.5. CAM-related gene expression profiling via quantitative real-time PCR

To identify the C_3_ to CAM transition time points, we assessed the expression of 28 CAM-related genes using quantitative real-time PCR (Table S1). Total RNA was isolated from leaf tissue using a CTAB method (Doyle & Doyle, 1987), and the quality and quantity were analyzed using a NanoDrop 1000 spectrophotometer (Thermo Fischer Scientific, San Jose, CA, USA). Five micrograms of total RNA were treated with RNase-free DNase I (New England Biolabs, Ipswich, MA, USA) to remove genomic DNA. cDNA was synthesized using a ProtoScript^®^ First Strand cDNA Synthesis Kit (New England Biolabs, Ipswich, MA, USA) according to the manufacturer’s manual. The cDNA was diluted to a final concentration of 50 ng/μl, and 5 μl was used in a 20 μl PCR reaction with 1 μl of each 10 μM primer and 10 μl of VeriQuest SyBr with Fluorescein kit (Affymetrix, Santa Clara, CA, USA). For each reaction, three technical replicates were included for each biological replicate. All amplifications were carried out on a CFX96 Touch™ Real-Time PCR Detection System (Bio-Rad, Hercules, CA, USA). The product size was confirmed by 2% agarose gel electrophoresis and the specificity of the amplicon was confirmed by melting curve analysis. Data were analyzed and quantified with the Bio-Rad CFX Manager software. The transporter gene *PLASMA MEMBRANE INTRINSIC PROTEIN* 1;2 (*PIP1;2*) was used as an internal PCR standard, as its expression is constant during the diurnal cycle in leaves from control and salt-treated *M. crystallinum* (Vera-Estrella *et al*. 2012). The relative expression levels were calculated by the 2^−ΔΔ C^_T_ method (Livak & Schmittgen 2001).

### 2.6. PEPC enzyme activity

For PEPC enzyme activity assays, 50 mg frozen leaf tissue was ground in liquid nitrogen to a fine powder, and extracted with 0.5 ml enzyme extraction buffer (200 mM HEPES-NaOH (pH 7.0), 5 mM dithiothreitol (DTT), 10 mM MgCl_2_ and 2% (w/v) PVP-40). The homogenized mixture was then centrifuged at 14,000 rpm, 4°C for 20 minutes and the supernatant was retained. To measure the activity of PEPC, 50 µl supernatant, 150 µl extraction buffer and 2.8 ml assay solution (100 mM HEPES-NaOH (pH 8.0), 5 mM DTT, 5 mM MgCl_2_, 10 mM NaHCO_3_, 2.5 mM phosphoenolpyruvete (PEP), 0.2 mM NADH and 0.0034 units/µl malate dehydrogenase (MDH)) were used. Absorbance at 340 nm was recorded for at least 5 min. PEPC enzyme activity was calculated as described previously (Chu *et al*. 1990).

### 2.7. Guard cell enrichment and construction of RNA-Seq libraries

For guard cell enrichment, parallel samples from control and salt-treated plants were collected at 12 am and 12 pm from day 5 to day 7 using a tape-peel method (Lawrence *et al*. 2018). Briefly, the abaxial epidermis was directly peeled off using Scotch Transparent tape, and adherent mesophyll cells were removed from the epidermis with a scalpel. After washing in basic solution (0.55 M sorbitol, 0.5 mM CaCl_2_, 0.5 mM MgCl_2_, 0.5 mM ascorbic acid, 10 μM KH_2_PO_4_, 5 mM 4-morpholineethanesulfonic acid (MES), pH 5.5 adjusted with 1 M KOH), the tapes with adherent epidermis were incubated with a cell wall digesting enzyme solution (0.7% cellulase R-10 (Yakult Honsha Co., Ltd, Tokyo, Japan), 0.025% macerozyme R-10 (Yakult Honsha Co., Ltd, Tokyo, Japan), 0.1% (w/v) polyvinylpyrrolidone-40 (Calbiochem, Billerica, Massachusetts, USA), and 0.25% (w/v) bovine serum albumin (Research Products International Corp., Mt Prospect, Illinois, USA) in 55% basic solution on a reciprocal shaker for 30 min at room temperature. Digested peels were washed three times with basic solution and quickly blotted dry on a filter paper, then immediately frozen in liquid nitrogen and stored at −80°C until RNA extraction. For each time point, three independent biological replicates were collected using three different sets of 120 plants. Total RNA was isolated using a CTAB method (Doyle & Doyle, 1987). After eliminating any genomic DNA contamination with RNase-free DNase I (New England Biolabs, Ipswich, MA, USA), mRNA was purified using a Dynabeads^®^ mRNA Purification Kit (Thermo Fischer Scientific, San Jose, CA, USA). RNA-seq libraries were constructed using the QuantSeq 3’ mRNA-Seq Library Prep Kit (Lexogen, Vienna, Austria) according to the manufacturer’s instructions. RNA sequencing was performed on an Illumina NextSeq 500 with 75 bp single end reads at the NextGen DNA Sequencing Core of the University of Florida, Gainesville, FL.

### 2.8. RNA-Seq data analysis

For gene expression quantification, a reference transcriptome was constructed using sequences from ice plant bladder cell (www.lsugenomics.org/data-from-our-group) (Oh *et al*. 2015) and root (dandelion.liveholonics.com/pothos/Mcr/data/reference/Mcr.transcript.fasta) (Tsukagoshi *et al*. 2015), sequences downloaded from NCBI, and Expressed Sequence Tags (ESTs) previously published (Cushman *et al*. 2008). Sequences from each reference were concatenated into a single fasta file to serve as a reference transcriptome. Because the *de novo* assemblies and transcripts were accessed from multiple sources, it was not known which transcripts were identical, or alternatively spliced isoforms, due to lack of a reference ice plant genome. To address the redundancy of the sequences in the reference and to collapse putative isoforms into a single transcript, Cap3 (Huang & Madan 1999) was used with default parameters (-a 20 –b 20 –c 12 –d 200 –e 30 –f 20 -g 6 –h 20 –i 40 –j 80 –k 1 –m 2 –n −5 –o 40 –p 90 –r 1 –s 900 –t 300 –u 3 –v 2 –w NA –x cap –y 100 –z 3). Assembled contigs representing putative collapsed transcript isoforms and singlet transcripts that were not homologous to other transcripts were kept as transcript references for alignments. These transcripts represent putative gene sequences. To exclude multiple isoforms of the same gene, the single longest contig was used for each gene. A single transcript representing each gene was used because downstream analysis of differential transcript expression was conducted at the gene level rather than the isoform level, as 3’ end sequencing or short read sequencing in general is not suitable for isoform level quantification. Low quality bases/reads were removed from the sequence data with Trimmomatic (Bolger *et al*. 2014) with parameters HEADCROP:0 LEADING:3 TRAILING:3 SLIDING-WINDOW:4:20 MINLEN:18. Reads were mapped to our reference transcriptome with RSEM (Li & Dewey 2011) version 1.2.31. Gene expression was measured as the number of reads that aligned to a given transcript (counts). Gene counts for each sample were consolidated into a matrix and imported into EdgeR (McCarthy *et al*. 2012) to conduct differential expression analysis. Differentially expressed (DE) transcripts were identified by comparing control versus salt treatment groups or day versus night groups from day 5, day 6, day 7, night 5, night 6 and night 7 sampling groups using a 2-fold change and an adjusted *p*-value threshold of 0.05. The RNA-Seq data have been deposited at the National Center for Biological Information (NCBI) Sequence Read Archive (SRA) under the accession number SRX3878746.

## 3. Results

### 3.1. Determination of the C_3_ to CAM transition timing

#### 3.1.1. Titratable acidity

Because of the nocturnal CO_2_ assimilation in CAM plants, high levels of malate accumulate in vacuoles of mesophyll cells before dawn, which increases cellular acidity. Changes of leaf titratable acidity measured at the start (8 pm) and end (8 am) of the dark period were used as a measure of CAM induction in *M. crystallinum* from day 4 to day 14 after 500 mM salt treatment. Salt-treated plants, measured from days 21 and 28, were used as positive controls because they use the CAM mode of photosynthesis only (Cushman *et al*. 2008; Vera-Estrella *et al*. 2012). A modest increase of overnight acidity was observed on day 6 in leaves of the salt-treated plants. The difference between treated and control plants became statistically significant on day 7, after which titratable acidity steadily increased (Figure 1A). It was noteworthy that there were no visible morphological differences between the control and salt-treated plants at day 7 (Figure 1B). Based on the nocturnal acid accumulation, we inferred that the C_3_ to CAM transition occurred between days 5 to 7 of salt treatment; thus, we collected samples from day 5 (no transition), day 6 and day 7 plants for follow-up analyses.

**Figure 1.**
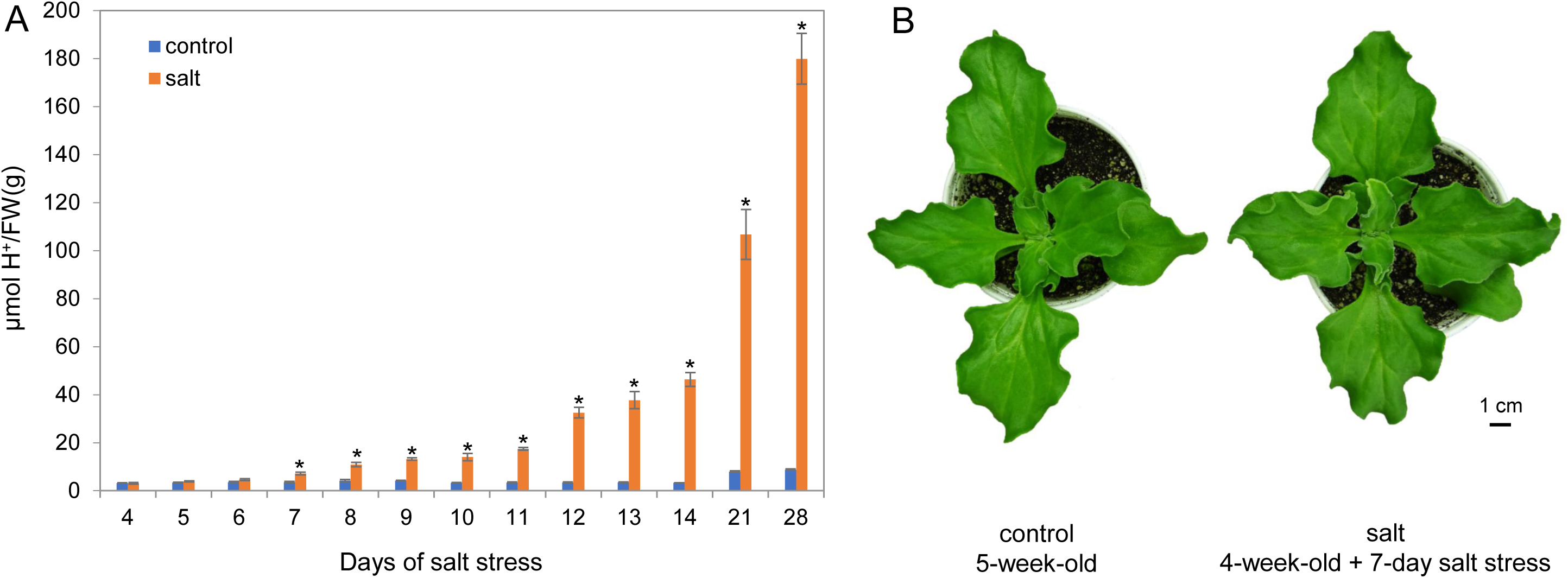
Leaf titratable acidity and morphological phenotype of *Mesembryanthemum crystallinum* under control and 500 mM NaCl treatment conditions. Four-week old plants were subjected to control (water) and 500 mM NaCl treatment for 4 to 28 days. (**A**) Measurement of leaf acidity was made at the start (8 am) of the photoperiod. Each bar represents mean of four replicates ± standard error (SE). Asterisks indicate statistical difference at *p*-value < 0.05 between the control and salt-treated plants as determined by t-tests. At the 7^th^ day, the *p*-value is 0.0062. (**B**) Images of control and 7-day salt-treated plants.

#### 3.1.2. Net CO_2_ exchange and stomatal movement

At day 5, control and salt-treated plants showed no difference in gas exchange parameters. However, day 6 salt-treated plants showed increased net CO_2_ uptake in the night and much lower net CO_2_ uptake during the day, compared to control plants (Figure 2A). Salt-treated plants from day 7 showed decreased net CO_2_ uptake during the day and positive net CO_2_ uptake in the night, significantly different from control plants (Figure 2A).

**Figure 2.**
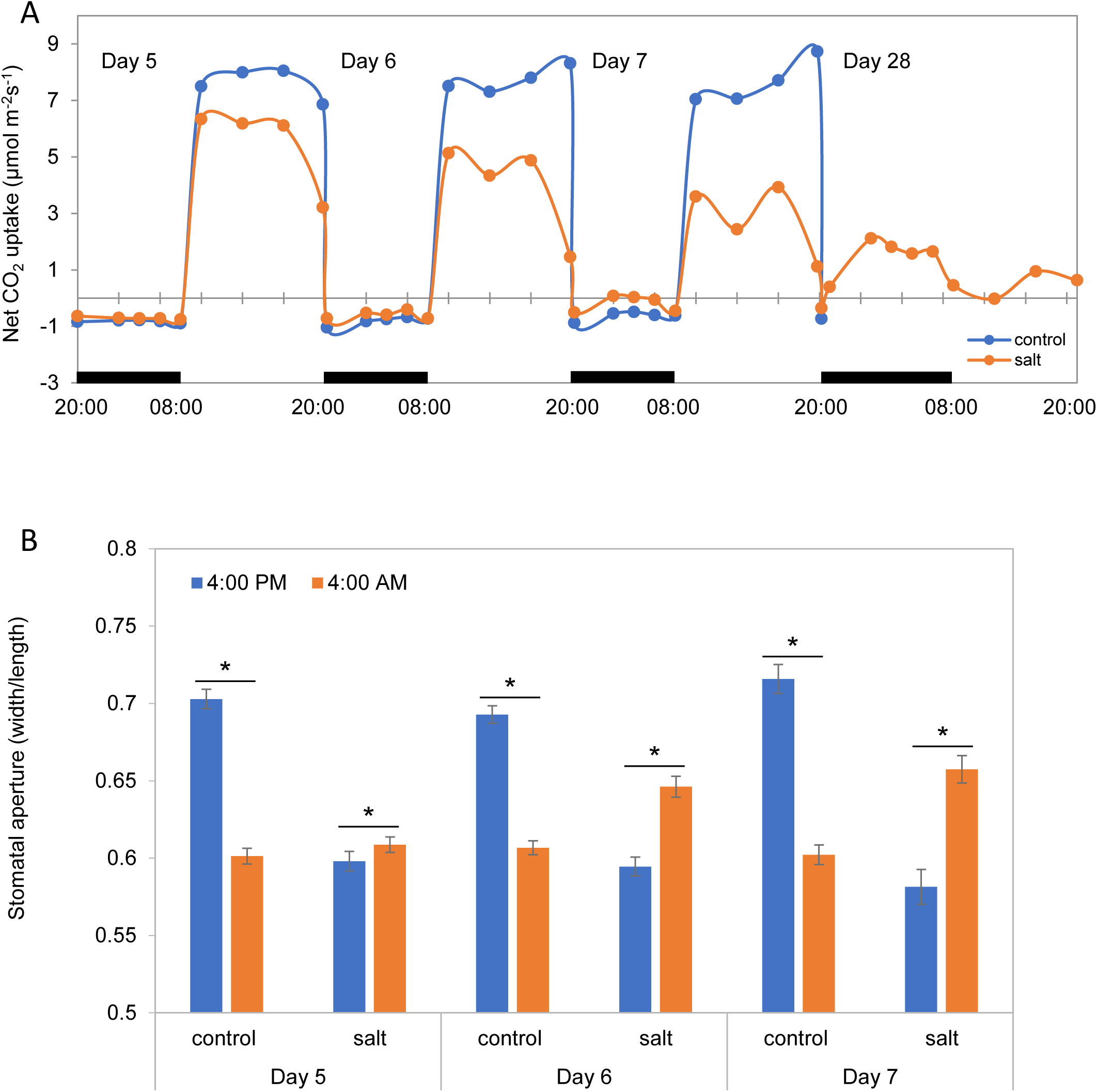
Day/night profiles of leaf net CO_2_ exchange and stomatal aperture in leaves of *M. crystallinum* grown under control (4-week old + 5 to 7 days with water) and salt treatment conditions (4-week old + 5 to 7 days with 500 mM NaCl). (**A**) Net CO_2_ exchange. The dark bars on the x-axis represent dark periods and each gas exchange profile is the average of three biological replicates. Day 28 CAM plants were included as positive controls. (**B**) Stomatal aperture of control and salt-treated plants. Data are mean ± SE of three independent experiments, each with 60 – 80 stomata for a total of at least 180 stomata. Asterisks indicate statistical difference at *p*-value < 0.05.

To further confirm the C_3_ to CAM transition time-points, stomatal aperture was measured in the control and salt-treated plants during the night (at 4 am) and the day (at 4 pm). These two time-points were chosen based on the result of net CO_2_ exchange (Figure 2A). As shown in Figure 2B, the control plants showed C3-type stomatal movement, i.e., they closed stomata during the night and opened stomata during the day. In contrast, the salt-treated plants showed an inversion in stomatal movement, starting on day 5. Stomatal aperture of the salt-treated plants increased consistently during the nights of day 6 and day 7 (Figure 2B). The gas exchange and stomatal movement results also support that the C_3_ to CAM transition of *M. crystallinum* occurs between day 5 to day 7 of the salt treatment.

#### 3.1.3. CAM-related gene expression profiles

To further validate the C_3_ to CAM transition, CAM-related genes were selected for analysis based on previous studies (Brilhaus *et al*. 2016; Cushman *et al*. 2008; Vera-Estrella *et al*. 2012). In total, 28 genes were selected, including PEP carboxylase (*PEPC*, the enzyme responsible for dark CO_2_ fixation) (Osmond 1978), PEPC kinase (*PPCK1*) (Dittrich 1976), 4-alpha-glucanotransferase or disproportionating enzyme (*GTF1*, involved in starch degradation) (Cushman *et al*. 2008), circadian clock associated 1 (*CCA1*) (Abraham *et al*. 2016), phosphoglycerate kinase (*PGK3*) (Brilhaus *et al*. 2016; Cushman *et al*. 2008), and ABA insensitive 1 (*ABI1*, involved in guard cell movement) (Abraham *et al*. 2016). From day 5 to day 7, control and salt-treated leaves were collected at 2 am and 4 am because two core CAM genes, *PEPC* and *PPCK1*, were reported to exhibit the highest expression levels at these time-points (Dodd *et al*. 2002). As shown in Figure 3A, *PEPC* exhibited a > 5-fold increase in transcript abundance at the two time points of day 6 in salt-treated plants compared to control plants and kept increasing up to > 200-fold at day 7. *PPCK1* showed a similar expression profile as *PEPC. ABI1* and *GTF1* also showed higher transcript abundances at the two time points of day 6 in the salt-treated plants, and the levels kept increasing at day 7. *ABI1* is an important ABA signaling component. The increased *ABI1* transcript abundance may contribute to the development of CAM (Figure 2B). *CCA1*, a key regulator of circadian rhythm in plants, showed a significant increase at 4 am of day 7 in salt-treated samples relative to control plants. *PGK3* showed significant decreases at the two time points of day 7 in the salt-treated samples. All six genes exhibited similar expression patterns at day 7 in the salt-treated plants as those reported in previous studies (Brilhaus *et al*. 2016; Cushman *et al*. 2008). Most of other 22 genes analyzed showed similar expression patterns as the above six genes at day 7 in the salt-treated plants (Table S2).

**Figure 3.**
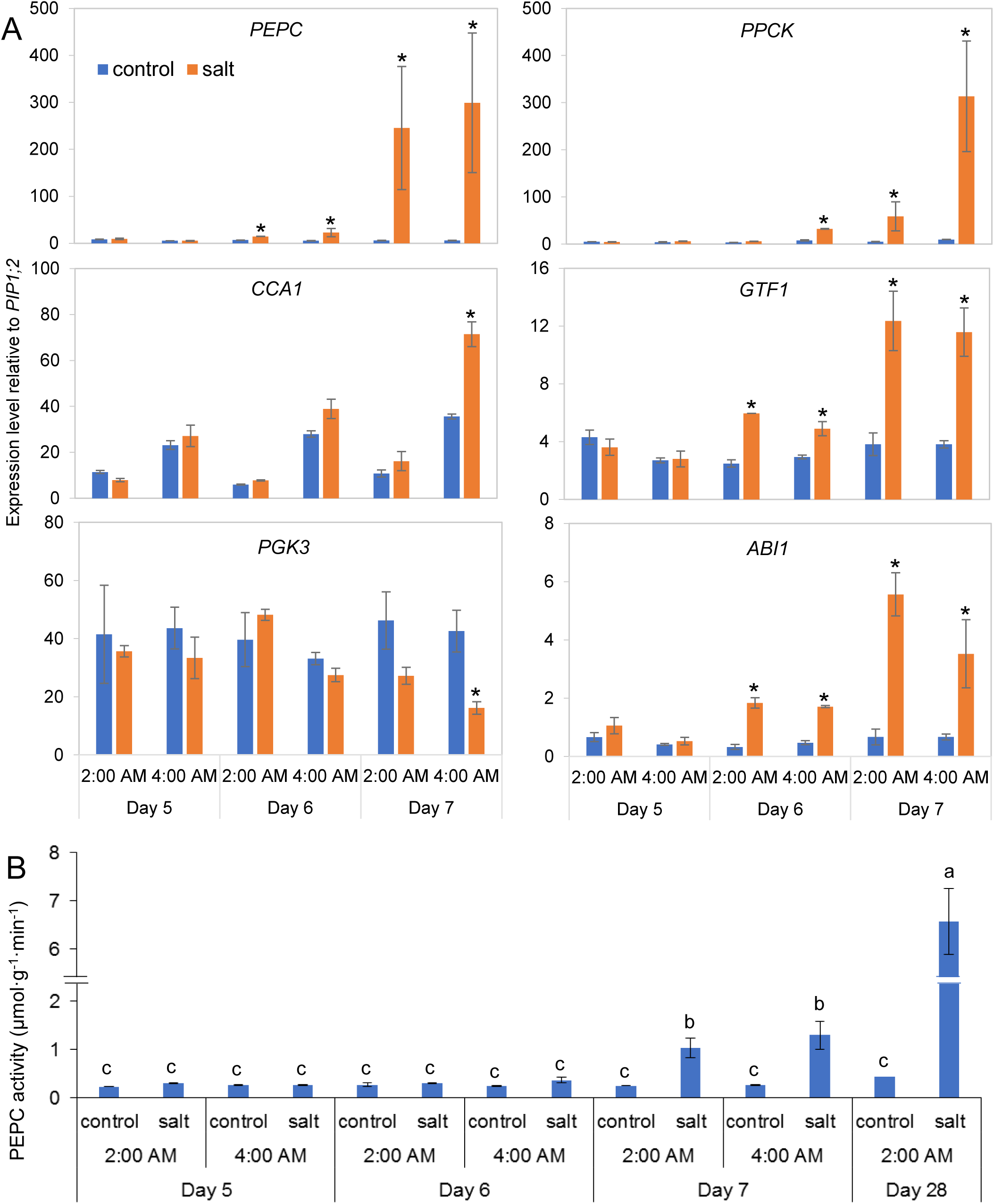
Expression profiles of CAM marker genes and PEPC activities during the C_3_ to CAM transition at 2 am and 4 am. (**A**) Transcript levels of *PEPC, PPCK, CCA1, GTF1, ABI1* and *PGK3* were determined by qRT-PCR. Error bars show the SE (n ≥ 3). Asterisks indicate a significant difference between control and salt-treated plants (Student’s *t*-test; *p-*value < 0.05). (**B**) PEPC enzyme activities were measured in three biological replicates. Bars represent standard deviation (SD) of the means. Two-way ANOVA and Tukey’s test were used for the PEPC activity analysis between different time points and different treated plants. Day 28 CAM plants were used as positive controls.

#### 3.1.4. PEPC enzyme activity

PEPC activity in the dark is much higher in CAM plants than in C_3_ plants (Chu *et al*. 1990). PEPC activity changes were measured in the samples collected at 2 am and 4 am of day 5 to day 7. As shown in Figure 3B, PEPC enzyme activity was significantly enhanced at the two day 7 time points in the salt-treated plants, as compared to the control. Taken together, the physiological data on titratable acidity, CO_2_ exchange, PEPC activity and stomatal movement, as well as the transcription of key CAM-related genes, demonstrate that the transition from C_3_ to CAM photosynthesis occurs between days 5 and 7.

### 3.2 Transcriptomics of *M. crystallinum* guard cells during the C_3_ to CAM transition

To investigate the regulatory mechanism(s) underlying the transition to inverse stomatal opening in *M. crystallinum*, we profiled the single-cell type transcriptome of guard cells using RNA-Seq during days 5, 6 and 7 at 12 am (night) and 12 pm (day). A total of 197,790,866 raw reads were acquired. Removal of low quality reads with Trimmomatic (Bolger *et al*. 2014) resulted in 188,147,736 high quality reads. By using previously published microarray and transcriptome data (Cushman *et al*. 2008; Oh *et al*. 2015; Tsukagoshi *et al*. 2015), we created a reference transcriptome, to which our short reads were mapped. In total, 43,165 different transcripts (including isoforms) were expressed in *M. crystallinum* guard cells (Counts Per Million (CPM) ≥ 10 in at least two biological replicates) (Table 1). Among these transcripts, 10,628 transcripts were not matched to the reference plant *Arabidopsis thaliana* transcripts (based on BLAST e-value ≤0.001, similarity ≥ 70).

**Table 1.**
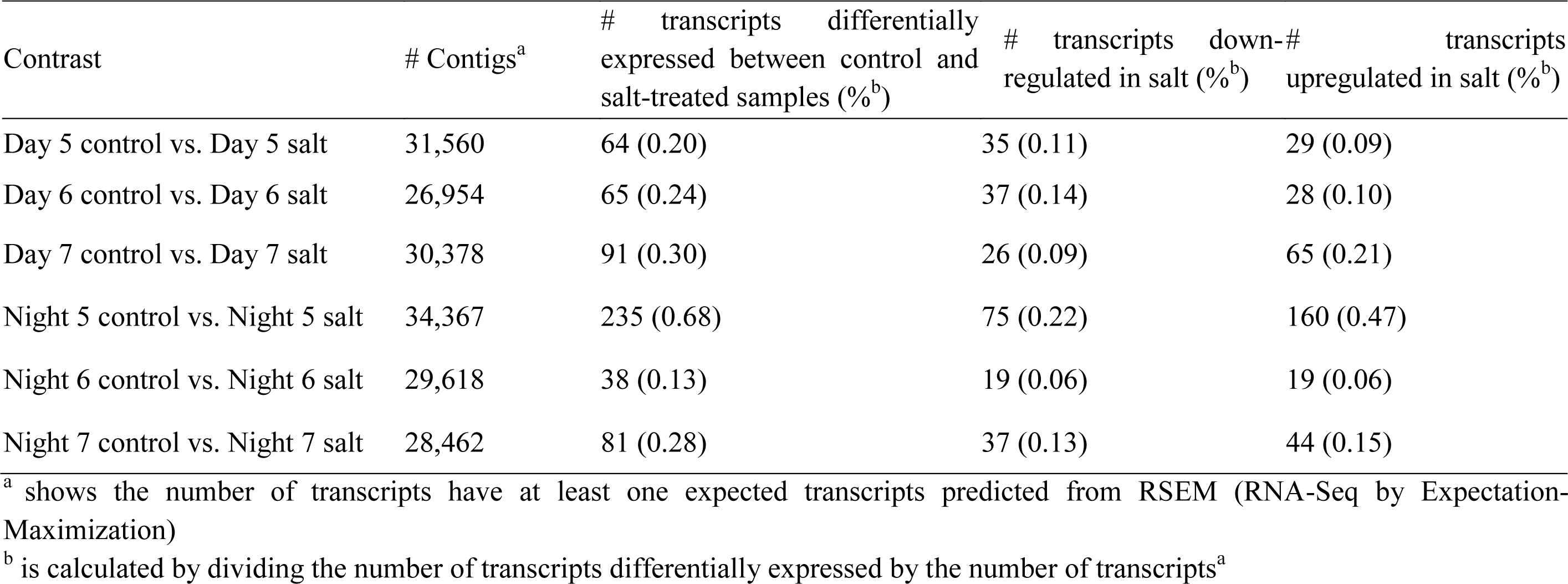
The number of differentially expressed transcripts in comparison of control and salt-treated samples

| Contrast | # Contigs<sup>a</sup> | # transcripts differentially expressed between control and salt-treated samples (%<sup>b</sup>) | # transcripts down-regulated in salt (%<sup>b</sup>) | # transcripts upregulated in salt (%<sup>b</sup>) |
| --- | --- | --- | --- | --- |
| Day 5 control vs. Day 5 salt | 31,560 | 64 (0.20) | 35 (0.11) | 29 (0.09) |
| Day 6 control vs. Day 6 salt | 26,954 | 65 (0.24) | 37 (0.14) | 28 (0.10) |
| Day 7 control vs. Day 7 salt | 30,378 | 91 (0.30) | 26 (0.09) | 65 (0.21) |
| Night 5 control vs. Night 5 salt | 34,367 | 235 (0.68) | 75 (0.22) | 160 (0.47) |
| Night 6 control vs. Night 6 salt | 29,618 | 38 (0.13) | 19 (0.06) | 19 (0.06) |
| Night 7 control vs. Night 7 salt | 28,462 | 81 (0.28) | 37 (0.13) | 44 (0.15) |

<sup>a</sup> shows the number of transcripts have at least one expected transcripts predicted from RSEM (RNA-Seq by Expectation-Maximization)
<sup>b</sup> is calculated by dividing the number of transcripts differentially expressed by the number of transcripts<sup>a</sup>

Significantly differentially expressed (DE) transcripts were defined as those with at least a 2-fold difference between control and salt-treated samples and an adjusted *p*-value < 0.05. A total of 495 transcripts showed significant changes at one or more time points by comparing the salt-treated and control plants; 369 of these DE transcripts have homologs in *Arabidopsis*. Among the 369 DE transcripts, *PEPC1* was found to increase at 12 am (night) of day 7 in the salt-treated samples (Table 2). This increased expression of *PEPC1* in guard cells correlates with our real-time PCR result in leaves (Figure 3).

**Table 2.**
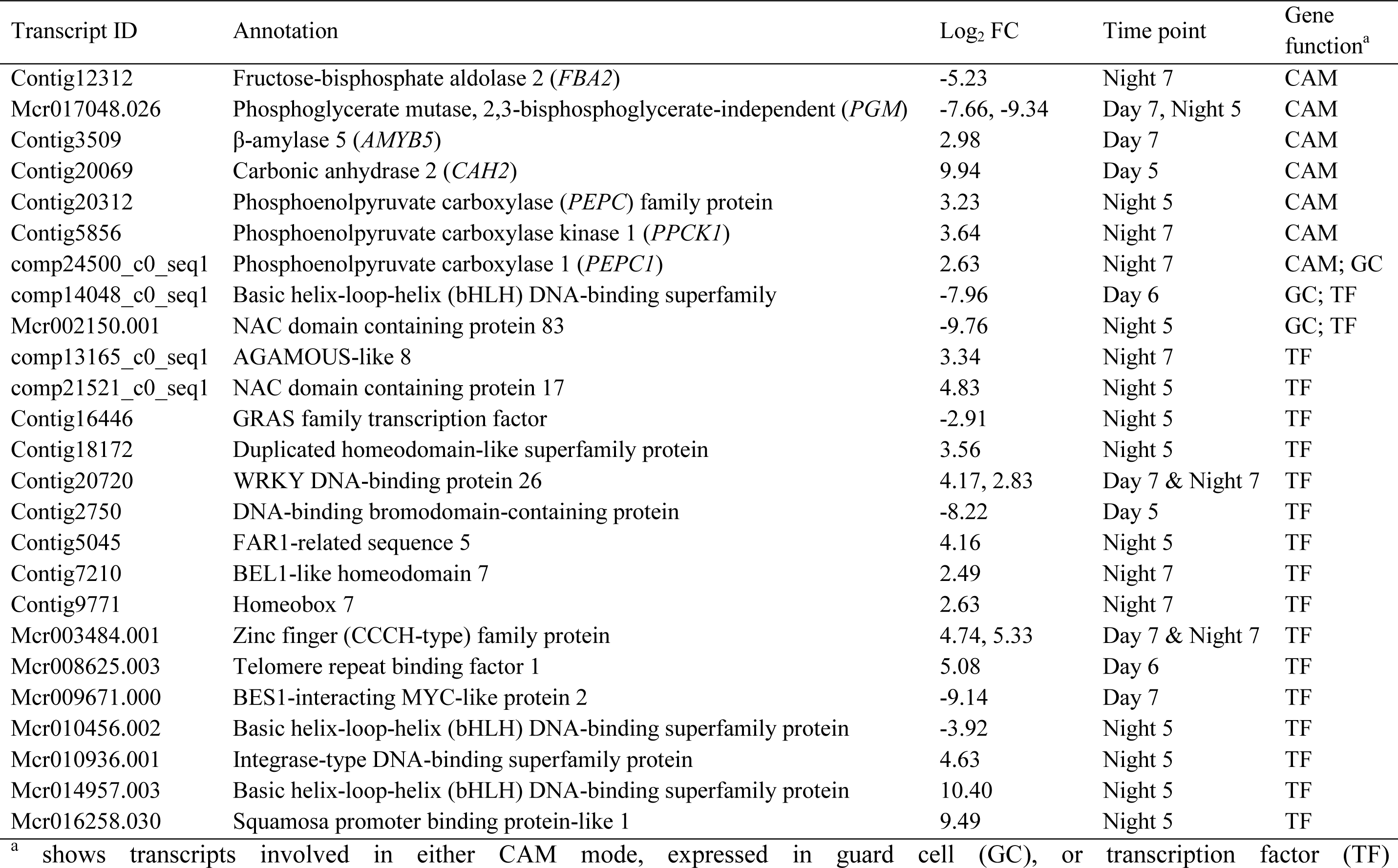
Functional categorization of selected transcripts differentially expressed in guard cells in response to salt stress. Significant change is expressed as a Log2 value of fold change of salt/control samples. For the full list of transcripts, please refer to Table S4.

Gene Ontology (GO) enrichment analysis of the 369 DE transcripts was performed using R package “clusterProfiler” (DOI: 10.18129/B9.bioc.clusterProfiler). The DE transcripts were enriched for the category “response to stimulus”, especially “response to abiotic stimulus” and “response to stress”. In terms of molecular function, the encoded proteins were enriched in catalytic activity (Figure 4). It is interesting to note that there was little overlap between the DE transcripts in the day samples (Figure 5A) or night samples (Figure 5B) at days 5, 6 and 7, suggesting different changes took place in the course of the early stages of transition from C_3_ to CAM. When the DE genes from day and night samples were compared, only about 10% of the DE genes were shared (Figure 5B). A small number of genes showed opposite change patterns under the day versus the night conditions (Table S3). For example, a papain-like cysteine protease showed high expression in control samples collected at night in day 5. At days 6 and 7, its expression levels started to decrease in control night and significantly increase in the day of salt-treated samples (Table S3). How this cysteine protease play a role in the CAM transition is not known. Nevertheless, this result suggests that the GC transcriptome is diurnally regulated during development of CAM (as are stomatal apertures).

**Figure 4.**
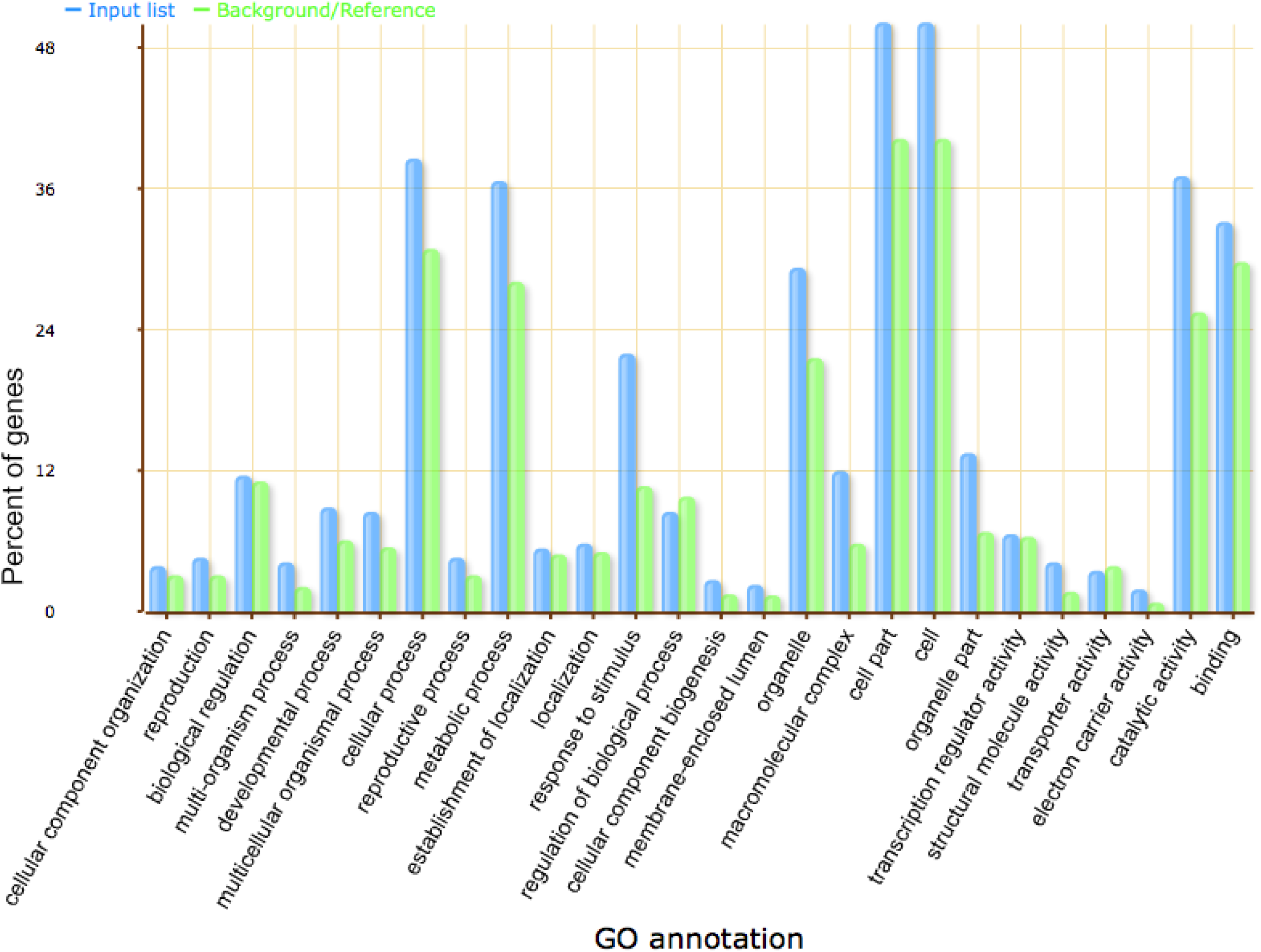
“Biological Process” functional Gene Ontology (GO) term classifications of DEGs. Please refer to Table S4 for detailed information of the DEGs.

**Figure 5.**
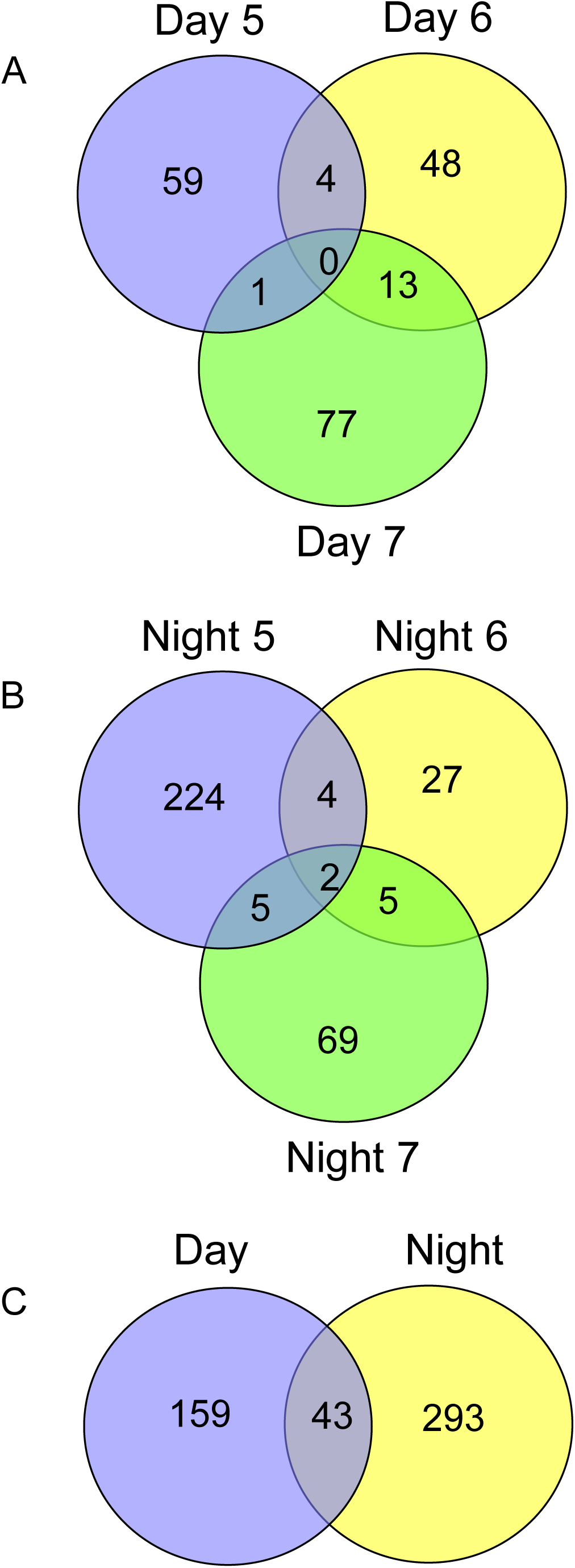
Overlap of DEGs identified at different time points. (A) Overlap of DEGs from different day time points. (B) Overlap of DEGs from different night time points. (C) Overlap of DEGs from day and night.

### 3.3 Abundance changes of previously identified guard cell transcripts during the C_3_ to CAM transition

To identify the CAM-related DE transcripts in our study, we compared these 369 DE transcripts with Cushman et al. (2008)’s microarray data, which identified 56 CAM-related genes and non-CAM isogenes in *M. crystallinum* that displayed maximal inverse expression between CAM-performing and C_3_ leaves. Seven out of the 56 genes were retrieved from our DE transcripts, including *PEPC1* (comp24500_c0_seq1), *PPCK1* (Contig5856), *BETA-AMYLASE 5* (Contig3509, *AMYB5*), *CARBONIC ANHYDRASE 2* (Contig20069, *CAH2*), *FRUCTOSE-BISPHOSPHATE ALDOLASE 2* (Contig12312, *FBA2*), *PHOSPHOGLYCERATE MUTASE* (Mcr017048.026, *PGM*) as well as another *PHOSPHOENOLPYRUVATE CARBOXYLASE* family protein gene (Contig20312) (Table 2). Among these seven genes, only *PPCK1* and *AMYB5* were not found in previous guard cell studies (Table 2). *PPCK1, PEPC1*, as well as another *PEPC* family gene, were up-regulated in our day 7 night samples, similar to the real-time PCR result on leaf tissue (Figure 3). The up-regulation of *PGM, CAH2, FBA2* and *AMYB5* in guard cells (Table 2) was also found in *M. crystallinum* leaves in a previous study (Cushman et al., 2008). However, in day 7 samples, *PGM* encoding a phosphoglycerate mutase up-regulated in leaves of the CAM plants (Cushman et al., 2008), was down-regulated in guard cells (Table 2). Overall, this analysis corroborates that guard cells themselves are transitioning to CAM because they exhibit many of the transcriptomic changes expected of cells undergoing that process. Additionally, guard cells also have their own transcriptional program during the transition.

### 3.4 Transcription factor changes during the C_3_ to CAM transition

It has been reported that in response to internal or external environment changes, transcription factors (TFs) exhibit more rapid expression changes than the bulk of the regulated genes (Jiao et *al*. 2003). Thus, the expression profiles of TF genes may in some way reflect the subsequent transcription activities regulated by them. In total, 18 TFs were identified among the 369 DE transcripts, and 14 of them were observed in other guard cell studies (Table 2). Based on the annotation from the *Arabidopsis* homologs, four TFs (Mcr002150.001, Mcr008625.003, Contig18172 and Contig9771) were identified in previous studies as responsive to salt or water deprivation stress in leaves or shoots (Table S4) (Ding *et al*. 2013; Seo & Park 2011; Yanhui *et al*. 2006). Since CAM mode in *M. crystallinum* is induced by abiotic stresses, such as salt and drought stress, these TFs may be general stress response genes and the others may have important roles in the C_3_ to CAM transition.

To evaluate how gene expression changes during the transition process, we grouped genes with similar pattern of expression using k-means clustering. Eleven clusters were retrieved, of which cluster 5 contained the greatest number of transcripts (Figure 6). In cluster 5, several genes showed differential expression profiles in the day samples, but the majority of transcripts showed increases in expression during the day 5 night. Only one gene, *ARGININE/SERINE-RICH SPLICING FACTOR 35* (Mcr012474.005) showed decreased expression during the day 7 night. In cluster 10, 20 of 24 transcripts have *Arabidopsis* homologs, and 17 of them were present in other guard cell studies (Figure 6). In this cluster, there are no differences in any samples during the day time, while all of them showed increases during the day 7 night (Figure 6). This cluster contains two key players in the CAM mode, *PEPC1* and *PPCK1*. It also includes three TFs: *AGAMOUS-LIKE 8* (AT5G60910), *HOMEOBOX 7* (AT2G46680) as well as BEL1-LIKE *HOMEODOMAIN 7* (AT2G16400) (Table 2).

**Figure 6.**
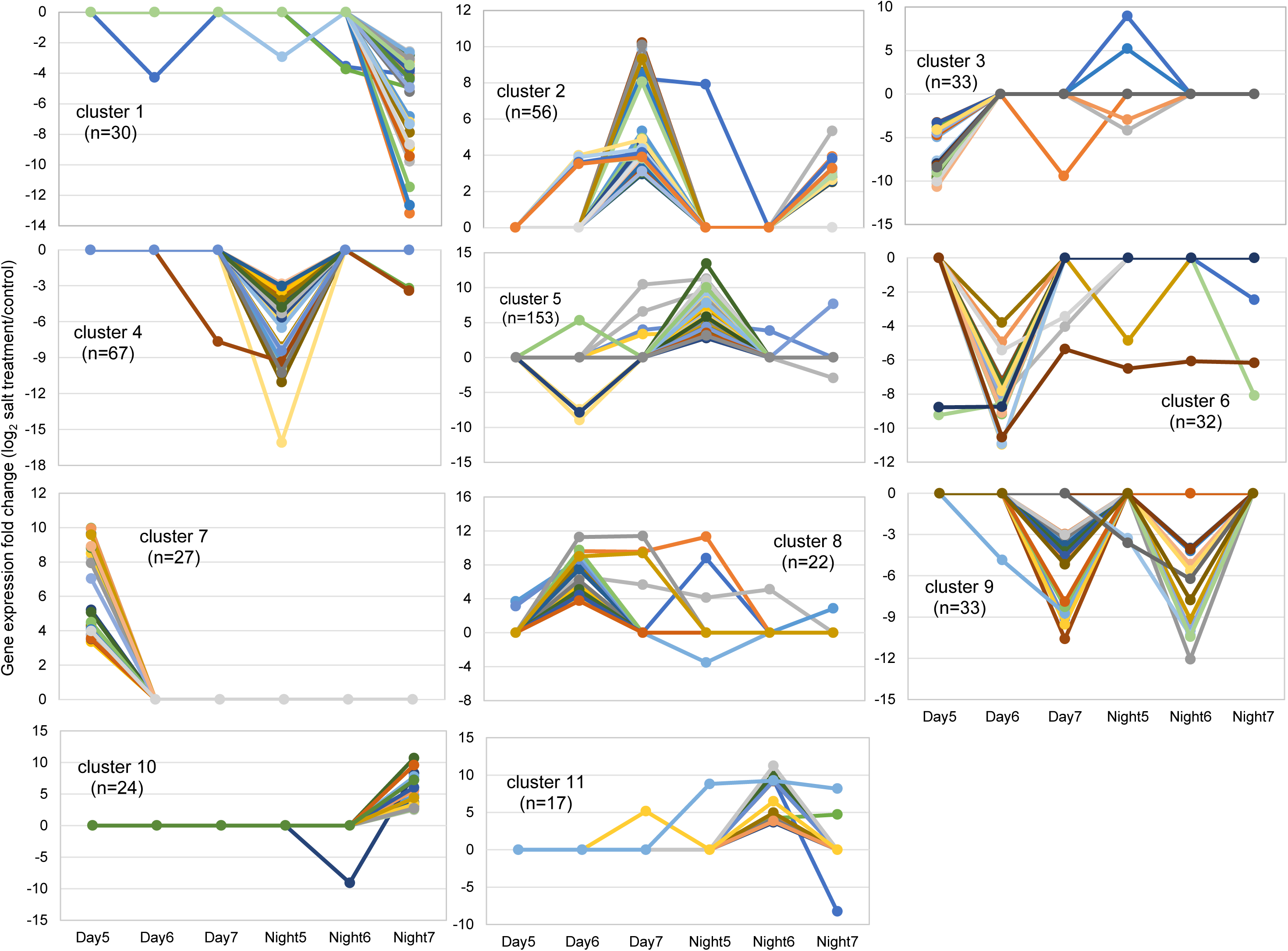
Clustering analysis of transcripts differentially expressed in guard cells of the C_3_ to CAM transition plants. *k*-means clustering algorithm (*k* = 8) were used. *n* indicates the number of transcripts in each cluster.

## Discussion

In this study, the C_3_ to CAM transition was induced by 500 mM salt treatment, and the transition time points were supported by several physiological parameters and molecular marker expression profiles (Figures 1-3). Furthermore, the cell-type specific transcriptome of guard cells was characterized during the C_3_ to CAM transition (Figures 4-6). Results are discussed here in light of the molecular mechanisms occurring in guard cells during the transition.

Although there have been many steady-state studies of C_3_ and/or CAM (e.g., Cushman *et al*. 2008; Davies & Griffiths 2012; Tsukagoshi *et al*. 2015), determination of the critical transition time-point was not reported before. The results of this study suggest that the C_3_ to CAM induction in *M. crystallinum* takes place within a short period of time (from day 5 to day 7), but that the full transition of photosynthesis to CAM is slow (another 2-4 weeks) (Figures 1, 2). Therefore, to determine the mechanisms underlying the C_3_ to CAM transition, we cannot rely on the analysis of one-time point. For the transcriptomic analysis of guard cells, we targeted six-time points from day 5 to day 7 of salt treatment.

We generated a reference transcriptome based on previously sequenced ESTs (Cushman *et al*. 2008), assembled transcripts (Oh *et al*. 2015; Tsukagoshi *et al*. 2015), and individual genes from *M. crystallinum* reported at NCBI. The genome size of *M. crystallinum* is 390 Mbp (Ha *et al*. 2014), with an estimate of 30,000 to 35,000 genes (De Rocher *et al*. 1990; Meyer *et al*. 1990). We constructed more than 180 thousand contigs, and identified 40,757 different transcripts (including isoforms) that were expressed in our samples. Among them, 10,628 unique transcripts in guard cells (no isoforms) were found to have homologs in Arabidopsis. Based on the previous studies of Arabidopsis guard cell transcriptomes (Bates *et al*. 2012; Bauer *et al*. 2013; Leonhardt *et al*. 2004; Pandey *et al*. 2010; Wang *et al*. 2011) and proteome (Zhao *et al*. 2008), about 30% of guard cell genes of *A. thaliana* had homologs expressed in *M. crystallinum* guard cells under our salt stress condition. From the RNA-Seq data, we identified 495 DE transcripts that showed significant changes at one or more time points of transition, of which 369 have homologs in Arabidopsis. Among these 369 DE transcripts, there were 199 up-regulated and 178 down-regulated transcripts in response to the salt treatment (Table 1). Both up- and down-regulated transcripts were enriched in “response to stress” and “cellular carbohydrate metabolic process”, suggesting a subset of genes involved in the two biological processes was employed in the course of transition. Key CAM molecular marker genes such as *PPCK1, PEPC1* and *GTF1*, as well as another *PEPC* family gene were up-regulated in the night 7 samples (Table 2, Table S4). This new observation suggests that guard cells switch from C_3_ to CAM; the timing of the differential expression in guard cells is also consistent with our physiological determination of the C_3_ to CAM transition time points (Figures 1-3). Another four CAM-related genes were also identified: *PGM, CAH2, FBA2* and *AMYB5*. Except for *PGM*, the other three genes showed similar expression profiles to those reported in leaves of CAM *M. crystallinum* (Table 2) (Cushman *et al*. 2008). *PGM*, which encodes a phosphoglycerate mutase was reported to be up-regulated in the plants performing CAM (Cushman *et al*. 2008), while in our study it was down-regulated in the day 7 samples. This may be due to difference in materials used since we targeted guard cells only, while Cushman and co-workers (2008) utilized the whole leaf in their study. Interestingly, PGM was shown to play an important role in guard cell functions. A null *pgm* mutant displayed defects in blue light-, abscisic acid-, and low CO_2_-regulated stomatal movements in Arabidopsis (Zhao & Assmann 2011). Another difference between our study and that of Cushman *et al*. (2008) is the time of sample collection: our samples were harvested during the C_3_ to CAM transition, while the previous study used leaves from plants performing complete CAM photosynthesis (Cushman *et al*. 2008). Therefore, this disparity indicates that the guard cells regulate transcription differently from those of leaves, either in response to salt stress or during transition and after transition to the CAM mode.

Notably, 18 TFs were identified among the 369 DE transcripts, and 14 of them were also detected in previous guard cell studies (Table 2). Among these 18 TFs, six of them were down-regulated, while 12 were up-regulated during the transition compared to the control guard cells from plants undergoing C_3_ photosynthesis. Four TFs (Mcr002150.001, Mcr008625.003, Contig18172, and Contig9771) may be related to salt or water deprivation stress response. These TFs are key players in the regulatory networks underlying plant responses to abiotic stresses and development processes (Coelho *et al*. 2018; Golldack *et al*. 2014; Hoang *et al*. 2017). Among them, *Mcr010456.002* (*At2g42280.1*), which encodes a basic helix-loop-helix (bHLH) TF, was implicated in stomatal movement through activation of genes encoding inwardly rectifying K^+^ channels (Takahashi *et al*. 2013). In addition, from the *k*-means clustering result (Figure 4), 26 DE transcripts (including the two CAM genes, *PEPC1* and *PPCK1*, present in cluster 10) were up-regulated in samples collected during the night of day 7 (Table S4), suggesting these genes were induced in the initial CAM guard cells. The significantly changed genes (including TFs) that are co-regulated with known CAM genes (e.g., *PPCK1* and *PEPC1*) can be expected to play important roles in the C_3_ to CAM transition process. In ice plants performing C_3_ photosynthesis, light increases leaf conductance and also promotes stomatal opening in isolated epidermal peels, while in plants performing CAM, stomatal opening in epidermal peels becomes unresponsive to light (Figure S1). This result and the RNA-Seq data from the isolated stomatal guard cells corroborate previous studies in facultative CAM species (Lee & Assmann 1992; Tallman *et al*. 1997), and demonstrate the presence in the guard cells themselves of molecular switches for the CAM inverse stomatal behavior, separate from mesophyll cells.

In summary, we induced CAM transition from C_3_ in *M. crystallinum* by 500 mM NaCl treatment and successfully determined the timing of the C_3_ to CAM transition. Furthermore, we characterized the guard cell transcriptomic changes during the critical transition process. The presence and the diel changes of CAM marker genes in stomatal guard cells indicate the guard cells themselves can transit from C_3_ to CAM. Many candidate genes (including TFs) were identified. Functional studies of these candidate genes in guard cells of either ice plant or the reference plant Arabidopsis are important future directions. In addition, these results indicate that efforts focused solely on engineering the mesophyll to introduce CAM into other species for improving WUE and stress tolerance may fail. Engineering both the mesophyll cells and guard cells is likely to be necessary.

## ACKNOWLEDGMENTS

We thank Christopher Krieg, Emily Sessa, Tianyi Ma, and Zepeng Yin for their help in obtaining data with the LI-6800, and Christopher Dervinis for his technical assistance in the RNA-Seq experiments. We also thank Dr. John Cushman from University of Nevada for providing the *Mesembryanthemum crystallinum* seeds and protocol for growing the plants. This work was supported by the University of Florida internal funds, with additional partial support from NSF Plant Genome Research Program Award #1444543, and NSF REU grant # 1560049 for the support to an undergraduate student Theresa M Kelley for carrying out part of this research.

## AUTHOR CONTRIBUTIONS

MK, SA and SC designed the research project. WK, MY and TK performed the experiments. MY and JN analyzed the transcriptomics data. JL helped with data interpretation and manuscript editing. MK and MY made the figures, and wrote the manuscript draft. All the authors read and approved the manuscript. SC finalized the manuscript for submission.

## Supporting Information

**Table S1.** List of primers used in this study.

**Table S2**. Expression analysis of CAM-related, circadian-related and guard cell signaling related genes in leaf samples using real-time RT-PCR.

**Table S3.** List of genes showing reversed expression patterns under control versus salt stress conditions in the guard cell RNA-seq dataset. The expression difference is expressed as log 2 value of fold change between day and night. Cells with “light orange” and “light blue” colors represent up-regulated in day and night samples at adjusted p-value < 0.05, respectively

**Table S4.** Functional categorization of 495 transcripts significantly differentially expressed in guard cells in response to salt stress. Significantly change is expressed as a Log 2 value of fold change of salt/control samples.

**Figure S1.** Stomatal aperture in leaves of C_3_ (4-week + 21-day water control) and CAM mode ice plants (4-week + 21-day salt treatment) under light and dark conditions. (**A**) Representative images showing stomatal aperture; (**B**) Stomatal aperture in leaves of C_3_ and CAM plants under light and dark. Data are mean ± SE of three independent experiments (*n* = 3) with 60 – 80 stomata for each replicate (i.e., a total of at least180 stomata for each experiment). Two-way ANOVA and Tukey’s test were used for stomatal aperture analysis between the C_3_ and CAM plants. For the images and stomatal aperture measurement, the epidermal peels were directly obtained from the plants, followed measuring the the width and length of the stomata under microscope.

