## Supplemental Figure 1 for "Molecular changes in *Mesembryanthemum crystallinum* guard cells underlying the C_3_ to CAM transition"

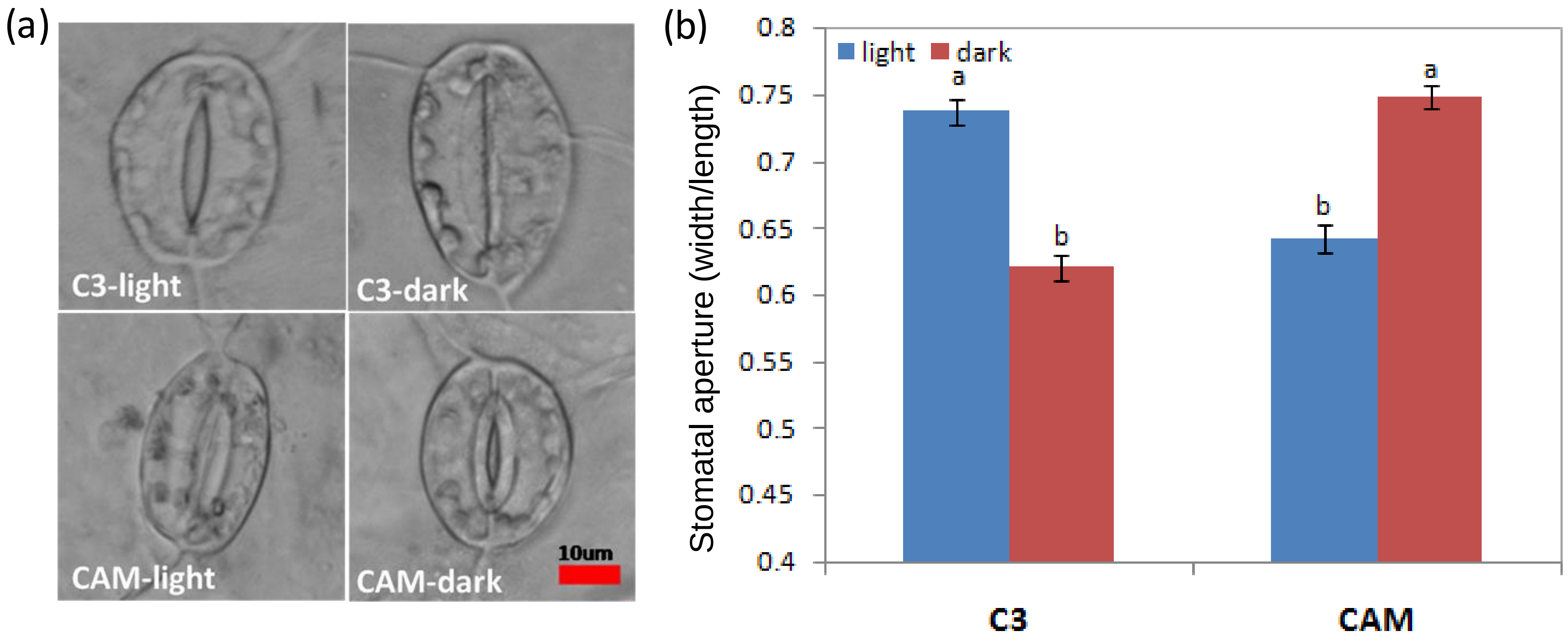


Figure S1. Stomatal aperture in leaves of C3 (4-week + 21-day water control) and CAM mode ice plants (4-week + 21-day salt treatment) under light and dark conditions. (**a**) Representative images showing stomatal aperture; (**b**) Stomatal aperture in leaves of C3 and CAM plants under light and dark. Data are mean ± SE of three independent experiments (n = 3) with 60 – 80 stomata for each replicate (i.e., a total of at least180 stomata for each experiment). Two-way ANOVA and Tukey's test were used for stomatal aperture analysis between the C3 and CAM plants. For the images and stomatal aperture measurement, the epidermal peels were directly obtained from plants, then the width and length of the stomata were measured.
